## Supplemental Information for "eEF3 promotes late stages of tRNA translocation on the ribosome"

### FOR

#### SUPPLEMENTAL FIGURES

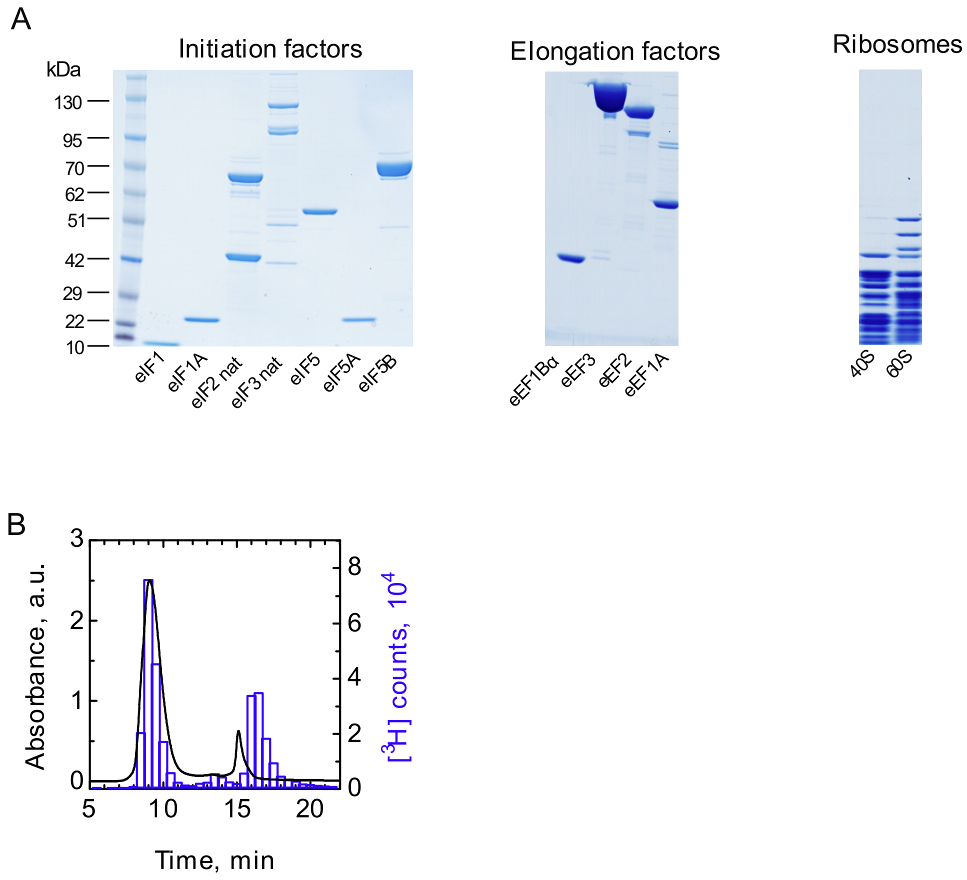


**Figure S1 Components of the reconstituted yeast translation system, related to Figure 1.** (**A**) Quality control of the initiation and elongation factors and the ribosomes by PAGE. For each component, 50 to 100 pmol were loaded on a NuPAGE 4 to 12% gradient gel. (**B**) Purification of the 80S IC by SEC on a Biosuite 450 using HPLC. The presence of 80S IC is identified as the overlap between the absorbance of the ribosomes at 260 nm (black) and the radioactivity from [$^3$H]Met-tRNA$_i^{Met}$ (blue).

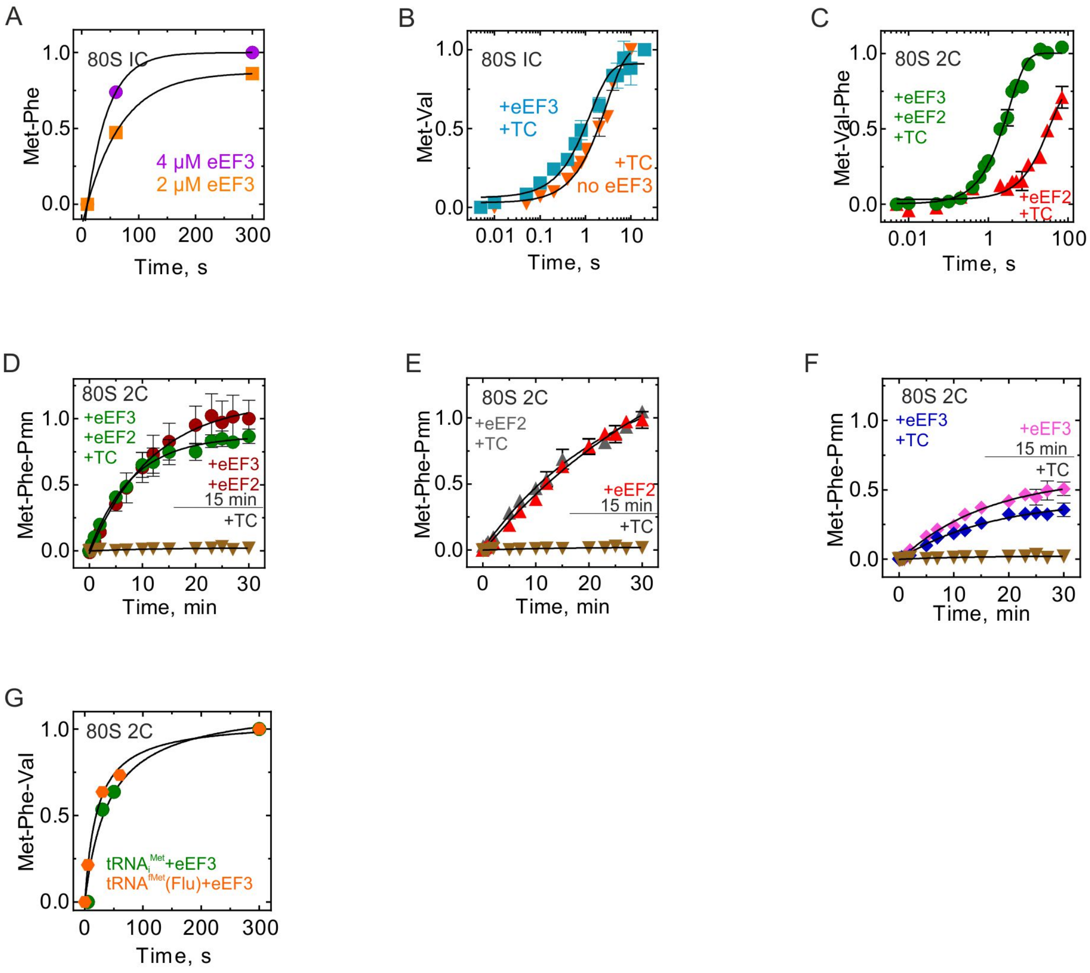


**Figure S2 Di- and tripeptide formation, related to Figure 2**. **(A)** Time courses of MetPhe formation at different eEF3 concentrations. **(B)** Met-Val formation monitored upon rapidly mixing initiation complexes (80S IC) with ternary complexes eEF1A–GTP–[$^{14}$C]Val-tRNA$^{Val}$ in the presence (cyan, 0.78±0.1 $s^{-1}$) or absence (orange, 0.34±0.03 $s^{-1}$) of eEF3 in a quench-flow apparatus, and the extent of peptide formation was analyzed by HPLC and radioactivity counting. Data presented as mean ± SEM of n = 3 biological replicates. **(C)** Met-Val-Phe formation upon rapid mixing of 80S complexes carrying MetVal-tRNA$^{Val}$ (80S 2C) with ternary complexes eEF1A–GTP–[$^{14}$C]Phe-tRNA$^{Phe}$ in the presence of eEF2 and eEF3 (green, 0.3±0.02 $s^{-1}$) or eEF2 (red, 0.03±0.006 $s^{-1}$). Data presented as mean ± SEM of n = 3 biological replicates. **(D-F)** Comparison of time courses of 80S 2C reaction with Pmn. 80S 2C complexes with MetPhe-tRNA$^{Phe}$ in the presence of eEF2 and eEF3 **(D)**, eEF2 **(E)** or eEF3 **(F)** with Pmn in a quench-flow apparatus. As indicated, the reaction was started either by mixing all components, or by addition of Pmn to a mixture of 80S 2C with the factors preincubated for 15 min. The extent of MetPhe-Pmn formation was analyzed by HPLC and radioactivity counting. Data presented as mean ± SEM of n = 3. For comparison, data from Figure 2B is plotted. **(G)** Met-Phe-Val formation upon rapid mixing of 80S complexes carrying either [$^{3}$H]Met-tRNA$_{i}^{Met}$ (green) or [$^{3}$H]Met-tRNA$^{fMet}$(flu) (orange) MetPhe-tRNA$^{Phe}$ (80S 2C) with ternary complexes eEF1A–GTP–[$^{14}$C]Val-tRNA$^{Val}$ in the presence of eEF2 and eEF3.

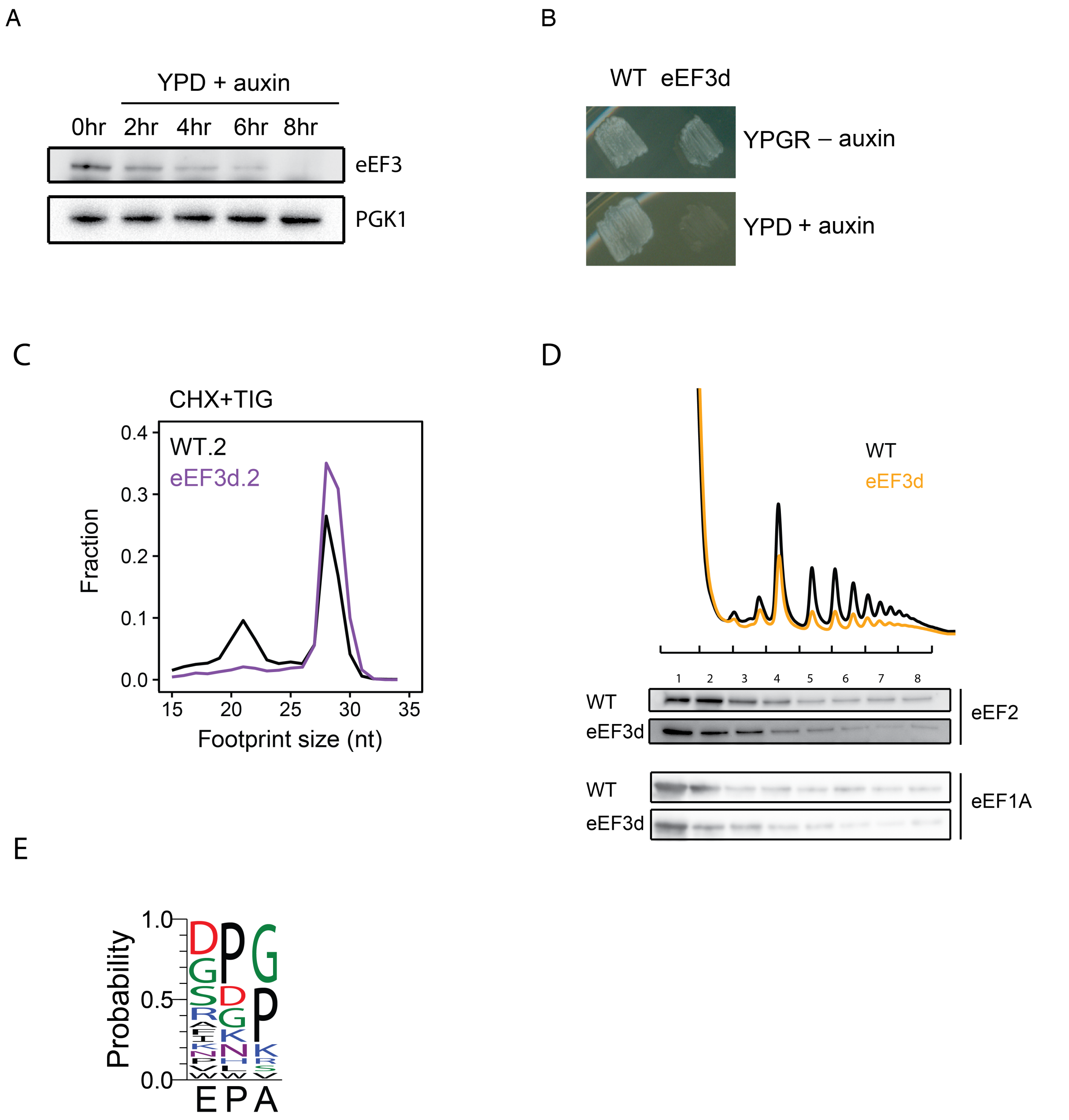


**Figure S3 Analysis of *in vivo* ribosome functional states by ribosome profiling, related to Figure 3.** (**A**) Immunoblot of eEF3 depletion over time. Same amount of cells were harvested at indicated time points, lysed and subjected to immunoblotting using antibodies against eEF3 or PGK1. (**B**) Growth of WT and eEF3d cells on YPGR and YPD+auxin plates. Plates were incubated at 30˚C for 1 days. (**C**) Size distribution of ribosome-protected footprints from biological replicates for Figure 3B. (**D**) Polysome profiles from WT or eEF3d cells (top). Fractions were analyzed by immunoblotting using antibodies against eEF1A or eEF2 (bottom). (**E**) Peptide motifs associated with ribosome pausing eEF3d cells, using motifs with a pause score greater than 4.

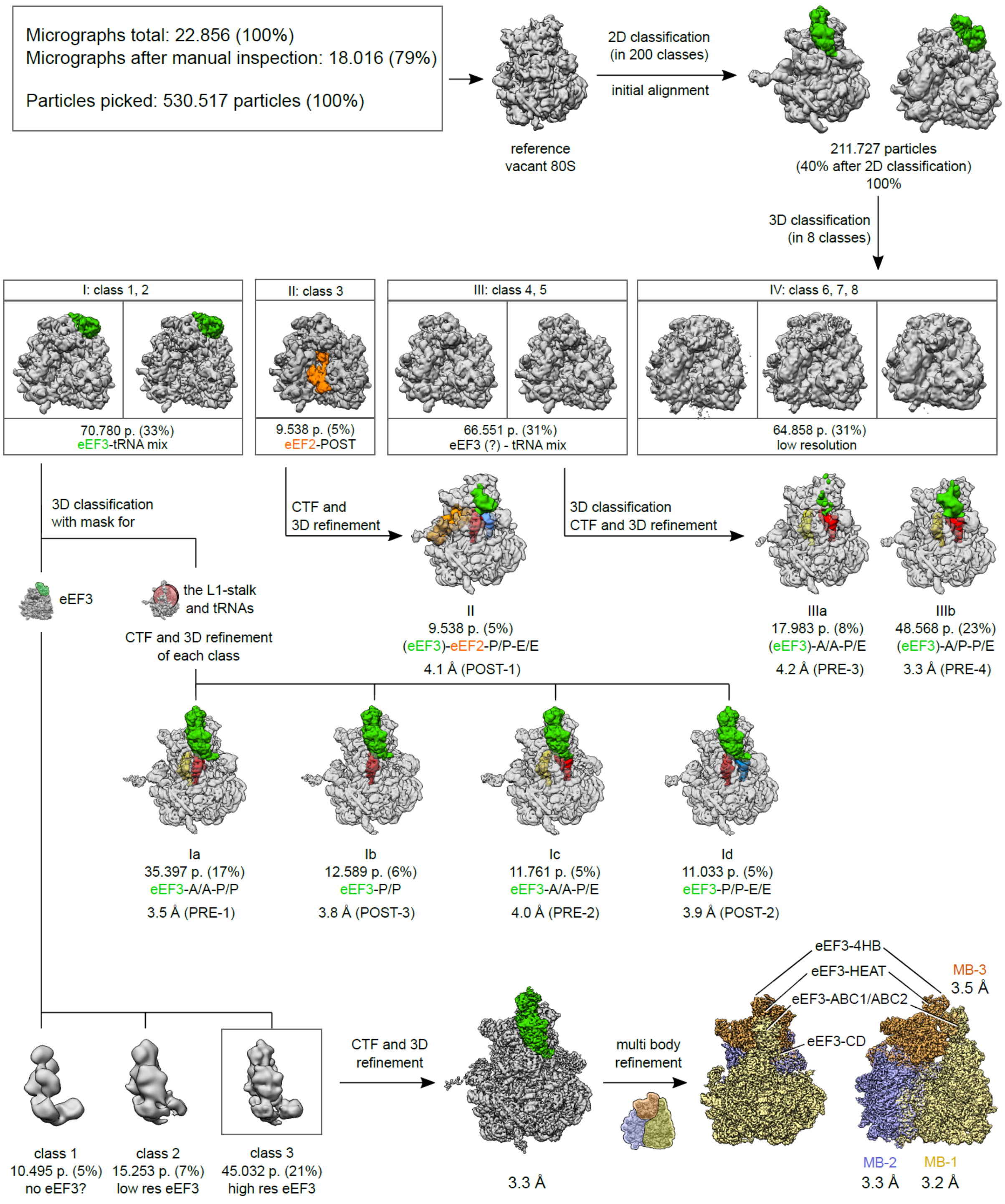


**Figure S4. 3D classification of the *S. cerevisiae* eEF3-80S complex, related to Figure 4.** Following 2D classification, 211,727 particles were initially aligned against a vacant *S. cerevisiae* 80S ribosome and subjected to 3D classification, sorting the particles into eight classes. The particles in class 1 and 2 bearing a stable eEF3-80S complex were joined in group I and subjected to two focused sortings using two different masks: the first encompassing the eEF3 ligand and the second covering the tRNAs, the L1-stalk and the eEF3-CD. The first classification allowed the sorting for a high resolution eEF3 bound volume (class 3, 21%, 45,032 particles), which was 3D and CTF refined resulting in a 3.3 Å final reconstitution. The final map was further multi body refined and provided a resolution of 3.2 Å for the LSU-eEF3 (ABC1/2, CD) (MB-1), 3.3 Å for the SSU body (MB-2) and 3.5 Å for the SSU head-eEF3 (HEAT, 4HB) (MB-3). The second sorting procedure enabled the classification of four EF3-bound non-rotated ribosomal classes with distinct tRNA occupancies (Ia-Id) with final resolutions denoted in the scheme. Class 3 (group II) showed a ribosomal species bearing eEF2, P/P- and E/E-tRNA and was finally refined to 4.1 Å. Class 4 and 5 (joined to group III) showed a rotated ribosome with mixed tRNA occupancy and after low pass filtering a disordered eEF3 ligand. This class was sorted further into two volumes (IIIa, IIIb) whereas each of them was 3D and CTF refined resulting in 4.2 Å and 3.8 Å resolution for the A/A- and P/E-tRNA occupied 80S (IIIa) and the A/P and P/E-tRNA bound ribsome (IIIb), respecively. Classes 6, 7, 8 (group IV) contained low resolution particles, which also showed a partial density for eEF3.

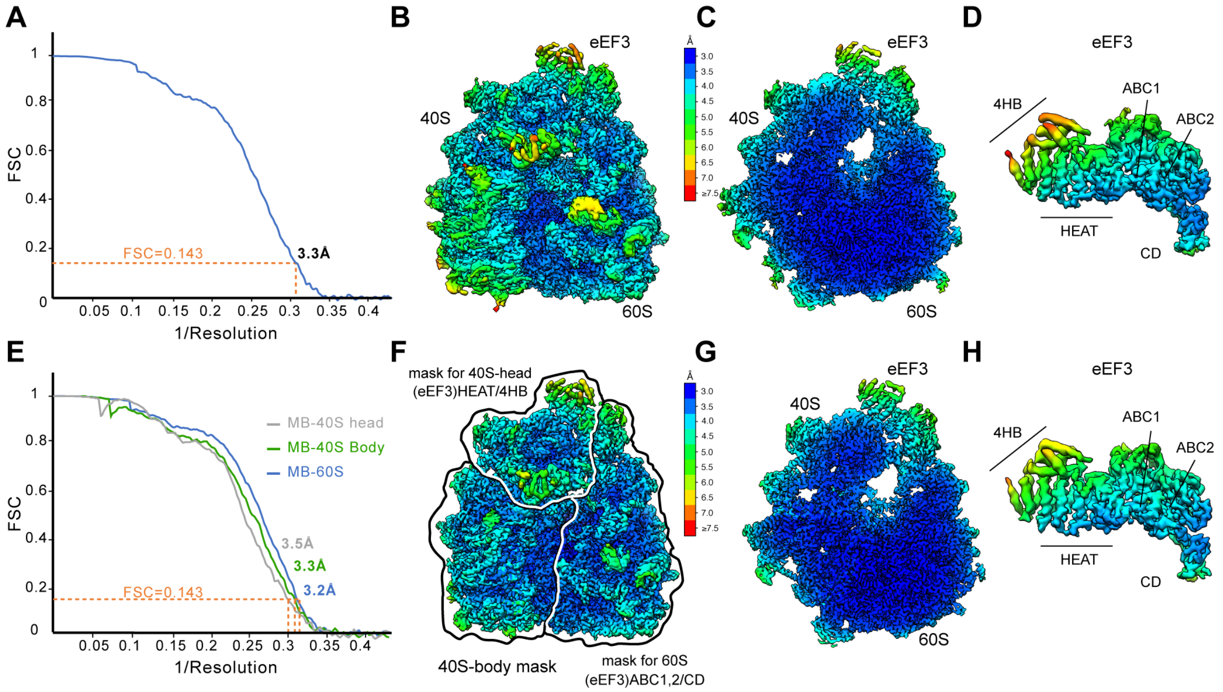


**Figure S5. Overview of the final refined cryo-EM reconstruction of the eEF3-80S complex, related to Figure 4.** (**A**) Fourier shell correlation (FSC) curve of the final refined cryo-EM map of the eEF3 ribosomal complex, indicating the average resolution of 3.3 Å, according to the gold-standard criterion (FSC=0.143). (**B**) Cryo-EM map of the 3.3 Å EF3-80S complex filtered and colored according to local resolution and (**C**) its transverse section. (**D**) View of the isolated density for eEF3 from (**B**). (**E**) FSC curve of the three multibody (MB) refined cryo-EM map of the eEF3-80S complex, indicating the average resolution of 3.2 Å for MB-60S, 3.3 Å for MB-40S body and 3.5 Å for MB-40S head. (**F**) Cryo-EM map of the MB refined EF3-80S complex filtered and colored according to local resolution. The outlines are depicting the three masks, which were used for the MB refinement. (**G**) Transverse section of the volume shown in (**F**). (**H**) Isolated density for eEF3 from (**F**). The scale bar is valid for all showed resolution panels.

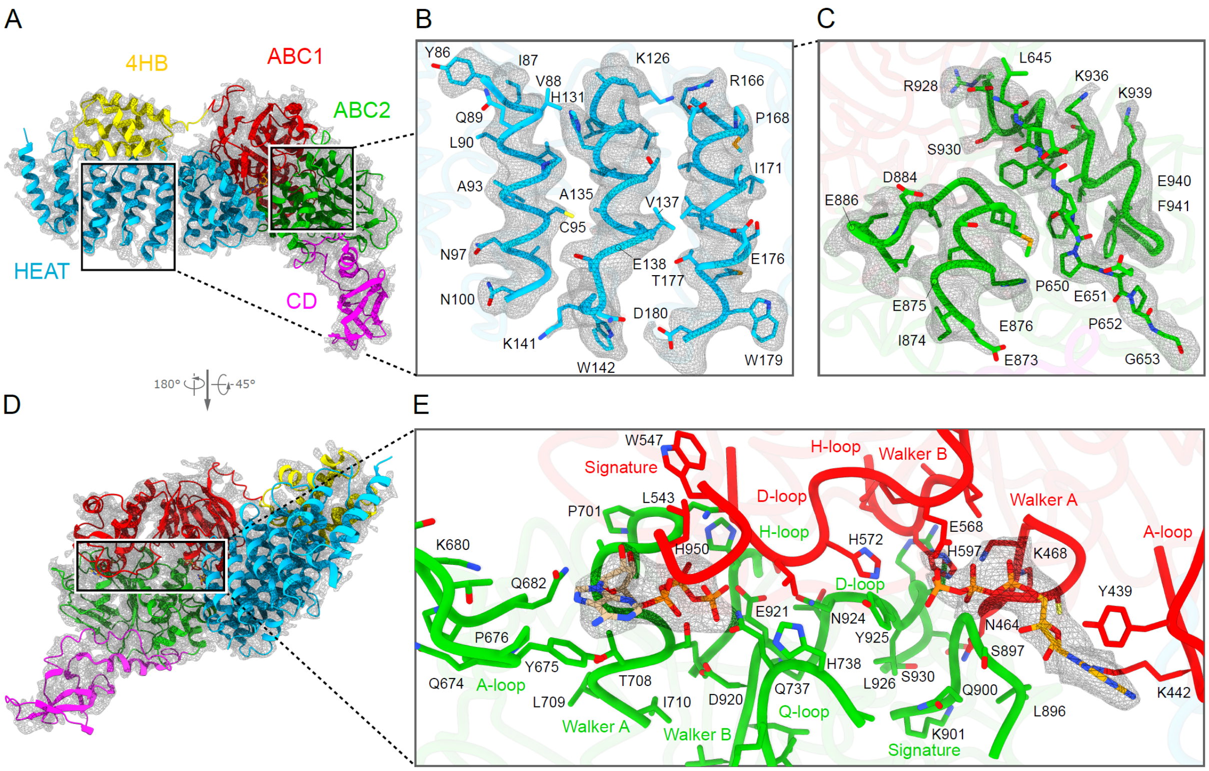


**Figure S6, related to Figure 4. Model for *S. cerevisiae* eEF3 with bound ATP molecules.** (**A**) Model of the 80S-bound eEF3 based on the multibody refined map (gray mesh) and colored by domain. HEAT (blue), 4HB (yellow), ABC1 (red), ABC2 (green) and CD (magenta). (**B**-**C**) Selected examples illustrating the quality of fit of the molecular model within (**B**) the HEAT repeat region and (**C**) the ABC2 domain to the unsegmented cryo-EM map (gray mesh). (**D**) eEF3-80S molecular model shown in (**A**), but in the orientation showing the ATP-binding cassettes and (**E**) zoom of the two nucleotide-binding sites formed by ABC1 and ABC2. For the bound ATP molecules cryo-EM map densities of the multibody refined map (dark grey mesh) are illustrated. The residues, which show density are shown as sticks and labelled.

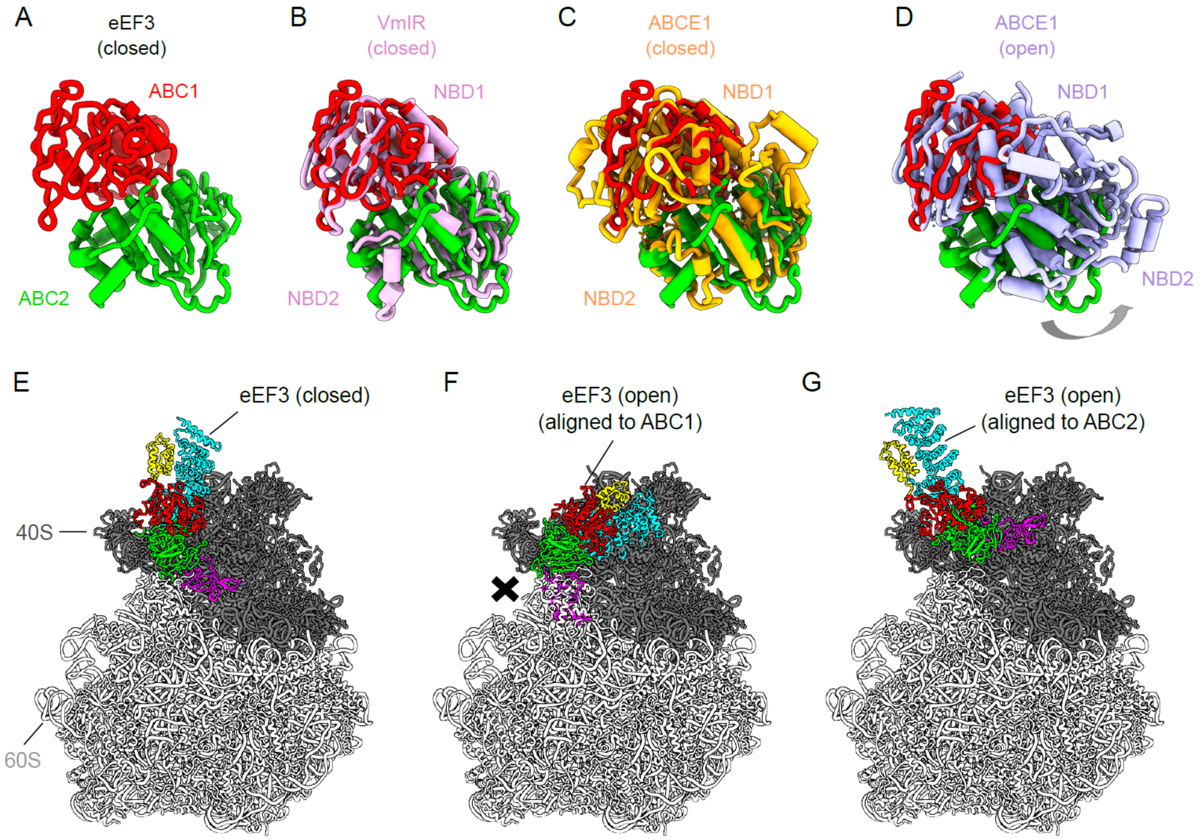


**Figure S7, related to Figure 4. The closed conformation of eEF3.** (**A**) The ABC1 (red) and ABC2 (green) domain of eEF3 in the eEF3-80S complex. (**B**-**D**) Conformation of the eEF3 NBDs with respect to other ABC proteins. Alignment (based on ABC1) of the eEF3-ABCs with (**B**) the closed conformation of the NBDs of the *B. subtilis* ABCF ATPase VmlR (pink, PDB: 6HA8) (Crowe-McAuliffe et al., 2018), (**C**) the closed conformation of *S. cerevisiae* ABCE1 (orange, PDB: 5LL6) (Heuer et al., 2017) and (**D**) the *E. coli* ABCE1 protein observed in the open conformation (violet, PDB ID: 3OZX) (Barthelme et al., 2011). (**E**) The eEF3 model in a closed conformation colored due to its different domain organization. (**F**-**G**) Incompatibility of eEF3 to the 80S ribosome in a potential opened conformation. The eEF3 model was aligned to (**F**) ABC1 or (**G**) ABC2 of the ABCE1 protein in an opened conformation (PDB ID: 3OZX) (Barthelme et al., 2011).

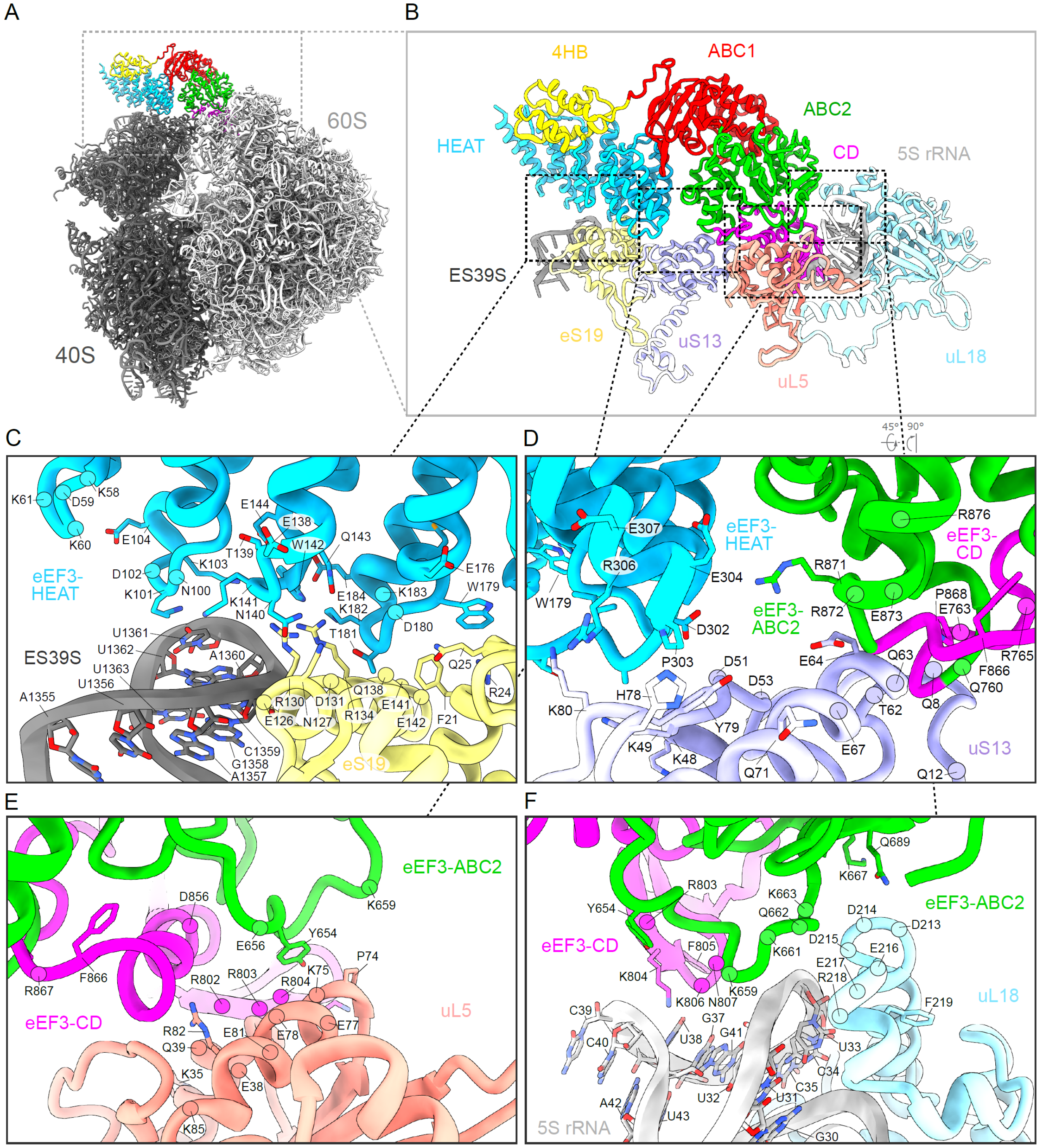


**Figure S8, related to Figure 4. Interactions of eEF3 with the 80S Ribosome.** (**A**) Overview of the eEF3-80S molecular model as well as (**B**) zoom on the eEF3 model colored due to its domain organization interacting with the proteins and rRNA of the LSU and the SSU. (**C**) Interactions of the eEF3-HEAT repeat region (blue) with ES39S of the 18S rRNA (dark gray) and eS19 (pale yellow) of the SSU. (**D**) The 40S protein uS13 (pastel violet) is forming bridging contacts with the HEAT repeats (blue) as well as the ABC2 (green) and the CD (magenta) of eEF3. (**E**-**F**) The eEF3-CD and the eEF3-ABC2 are interacting with (**E**) the LSU protein uL5 (coral) as well as with (**F**) the 5S rRNA (light gray) and uL18 (pale blue).

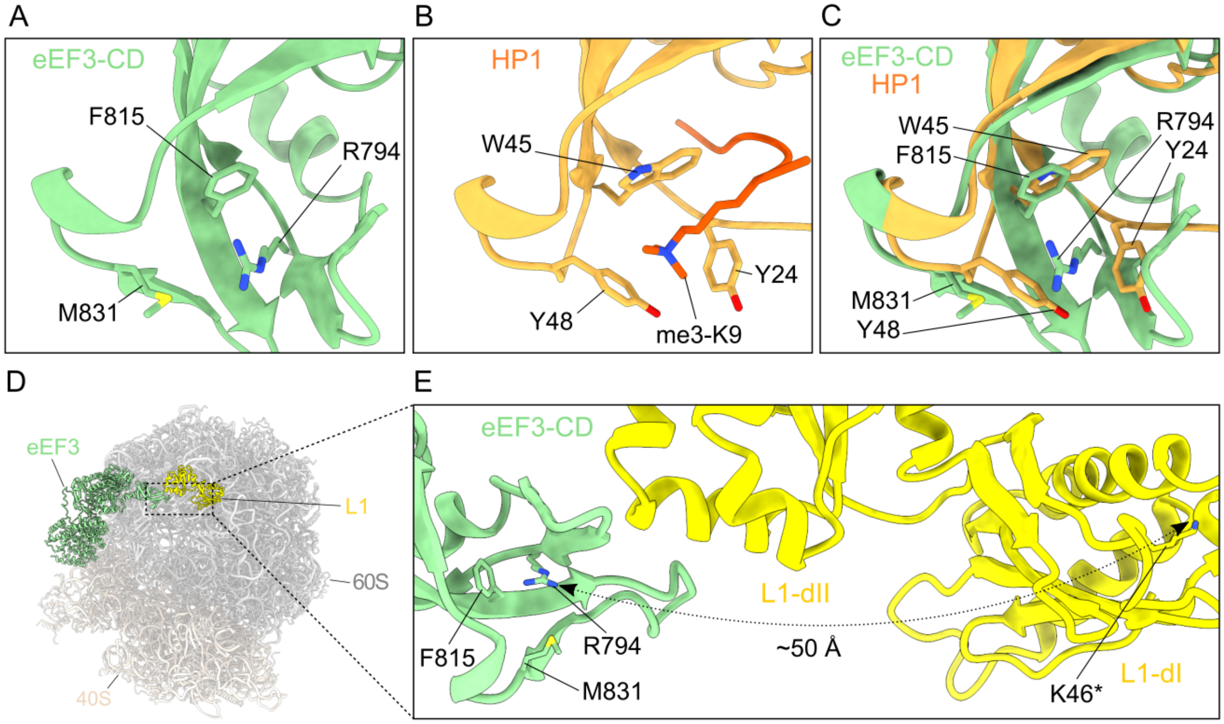


**Figure S9. The hydrophobic pocket of the eEF3 CD, related to Figure 5.** (**A**) The eEF3-CD hydrophobic pocket based on the alignment with (**B**) the *D. melanogaster* HP1-CD. (**C**) Overlay of the eEF3 and HP1 CD based on their sequence. (**D**) eEF3-80S molecular model highlighting eEF3 (pale green) and the L1 protein (yellow) and its (**E**) zoom showing the magnitude of the distance between the eEF3-CD hydrophobic pocket (based on the alignment with HP1 shown in (**C**)) and the K46 of the L1-dI. K46* labels the lysine, which is getting methylated by the Seven-β-strand methyltransferase (Webb et al., 2011).

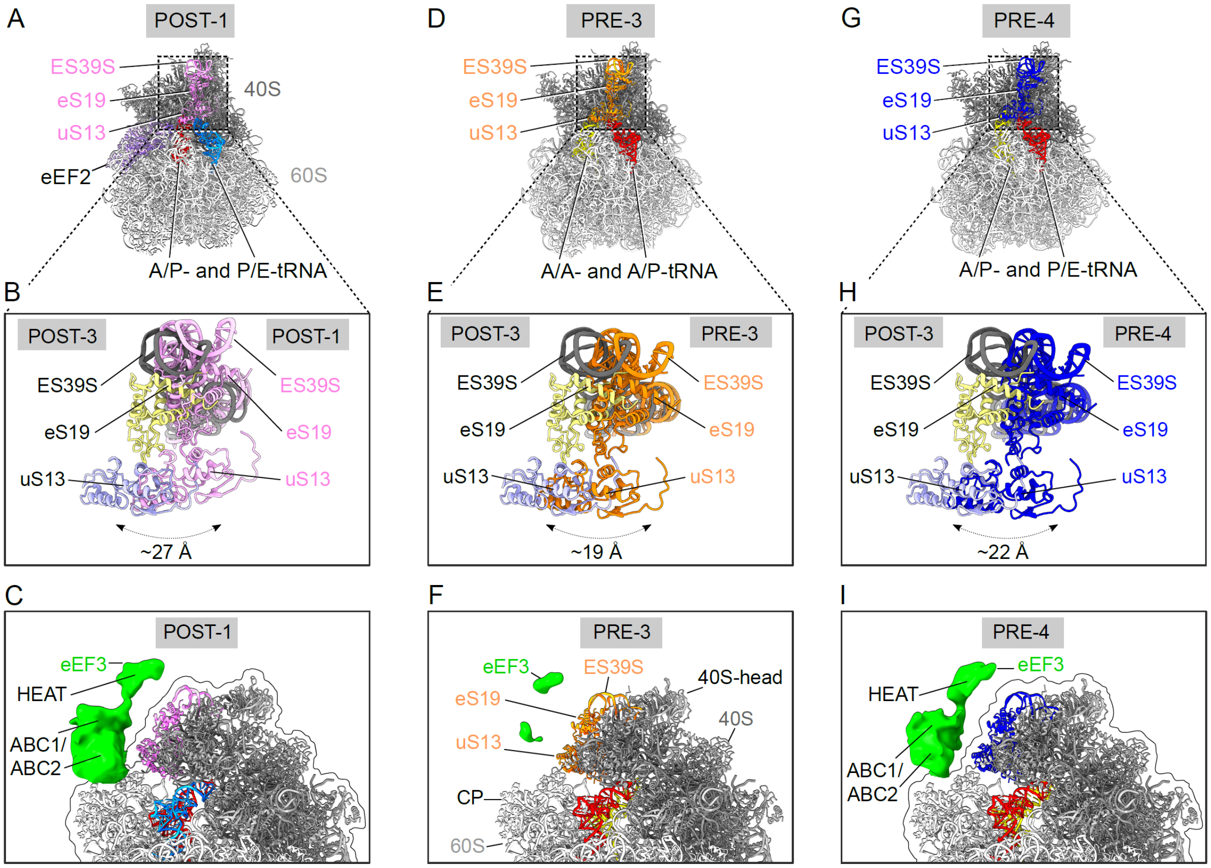


**Figure S10. The rotated 80S bound to disordered eEF3, related to Figure 6.** (**A**) Rotated 80S molecular model corresponding to the POST-1 state (disordered density for eEF3 is not shown). The ribosomal residues ES39S, eS19 and uS13 responsible for eEF3-HEAT binding are depicted in pink. (**B**) Zoomed view of ES39S, eS19 and uS13 highlighted in (**A**) overlaid with corresponding residues from the non-rotated POST-3 state. The arrows are showing the magnitude of the movement of these residues from a non-rotated POST-3 to a rotated POST-1 state in (**A**). (**C**) Segmented density for the disordered eEF3 from the POST-1 cryo-EM map highlighting the lack of association of the eEF3-HEAT repeat region with the moved ES39S, eS19 and uS13 resulting from the subunit rotation. (**D**) 80S model in rotated PRE-3 state depicting ES39S, eS19 and uS13 in orange and the (**E**) corresponding zoom of the displacement of the mentioned residues in the rotated ribosome. (**F**) Isolated density for the potential disordered ligand from the PRE-3 volume showing barely any density for eEF3. (**G**) PRE-4 ribosomal model with ES39S, eS19 and uS13 colored in blue as well as (**H**) the corresponding enlargement of these residues compared to the residues in the POST-3 state. (**I**) Segmented density of the eEF3 ligand from the rotated PRE-4 state. The isolated densities in (**C**), (**F**) and (**I**) were low pass filtered to 8 Å.
